# *β*-motifs and molecular flux promote amyloid nucleation at condensate interfaces

**DOI:** 10.64898/2026.04.09.717507

**Authors:** Subhadip Biswas, Davit A Potoyan

## Abstract

Biomolecular condensates are increasingly implicated as intermediates in the formation of pathological amyloid assemblies, yet the mechanisms by which sequence-encoded structural motifs and non-equilibrium molecular transport cooperate at condensate interfaces remain incompletely understood. Here, we introduce Flux-Driven Molecular Dynamics (FD-MD), a simulation framework that combines sequence-encoded *β*-prone interactions with sustained molecular influx to examine fibril formation at condensate interfaces. Within this framework, we establish three main results. First, a scaling analysis of orientational entropy suggests that condensate interfaces can enhance nucleation relative to the bulk by as much as two orders of magnitude, by reducing the entropic cost of coalignment of rigid *β*-prone segments. Second, varying segment rigidity and molecular supply rate organizes a non-equilibrium phase diagram with four interfacial growth morphologies, ranging from uniform wetting to fibrillar protrusions and inter-condensate bridging networks. Third, directional fibril elongation displays an inverse relationship with drift velocity, consistent with a mechanism in which higher transport rates to the interface favor planar saturation over directed tip incorporation. Together, these results support a picture in which condensate interfaces can act as kinetically favorable nucleation environments, sequence-encoded rigidity helps determine whether interfaces remain liquid-like or become fibrillar, and molecular flux emerges as an additional control axis in the model for condensate aging trajectories.

## I. INTRODUCTION

Biomolecular condensates are now recognized as central regulators of cellular organization, forming dynamic, membraneless compartments that coordinate biochemical reactions in space and time [1–4]. In their functional states, these assemblies exhibit liquid-like material properties, including rapid internal rearrangements and molecular exchange with the surrounding environment, enabling robust yet adaptable cellular responses [3, 5–9]. Accumulating evidence, however, demonstrates that biomolecular condensates can act as metastable intermediates along non-equilibrium pathways toward more heterogeneous and solid-like states. As condensates age, they may undergo liquid-to-solid transitions that culminate in the formation of *β*-rich fibrillar assemblies implicated in neurodegenerative disease [10–16]. Importantly, these transitions are not governed solely by bulk thermodynamics [17–19]. Instead, a growing body of experimental work indicates that condensate interfaces serve as preferential nucleation microenvironments, where reduced dimensionality, molecular crowding, and interfacial anisotropy lower kinetic barriers to fibril nucleation [20–29]. Across diverse systems, including hnRNPA1 and Tau low-complexity domains, as well as minimal ATP-based droplets, fibril formation consistently initiates at condensate interfaces and is sustained by continuous recruitment of molecules from the dilute phase [30–37]. Despite these advances, the kinetic mechanisms by which sequence-encoded structural motifs and driven molecular exchange cooperate at interfaces to promote fibrillization remain poorly understood.

Existing computational approaches face fundamental limitations in their ability to resolve these processes [38–40]. Atomistic simulations can capture sequence-specific interaction grammars but are restricted to system sizes and timescales far below those required to observe interface-mediated fibril growth. Conversely, coarse-grained models can access mesoscale condensate dynamics but typically lack explicit representations of *β*-prone structural motifs or sustained non-equilibrium molecular exchange. As a result, to our knowledge, existing simulation frameworks do not simultaneously integrate (i) sequence-encoded *β*-forming interactions, (ii) interfacial alignment and anisotropy, and (iii) persistent non-equilibrium influx, three ingredients increasingly recognized as essential for condensate-to-fibril transitions.

Here, we address this gap by introducing Flux-Driven Molecular Dynamics (FD-MD), a non-equilibrium simulation framework that captures the coupled structural and kinetic determinants of fibril growth at condensate interfaces (Fig. 1). FD-MD combines sequence-encoded rigid, *β*-prone segments with a controlled molecular influx that sustains interfacial recruitment beyond purely diffusive limits. Using the FD-MD framework, we find that rigid segments preferentially align at condensate surfaces to nucleate surface-bound protofilaments, which elongate into surface-anchored fibrillar protrusions through flux-sustained growth and progressively remodel condensate interfaces. Together, our results provide a mechanistic framework linking sequence-encoded structure, interfacial geometry, and driven mass transport to aging pathways in biomolecular condensates.

**FIG. 1.**
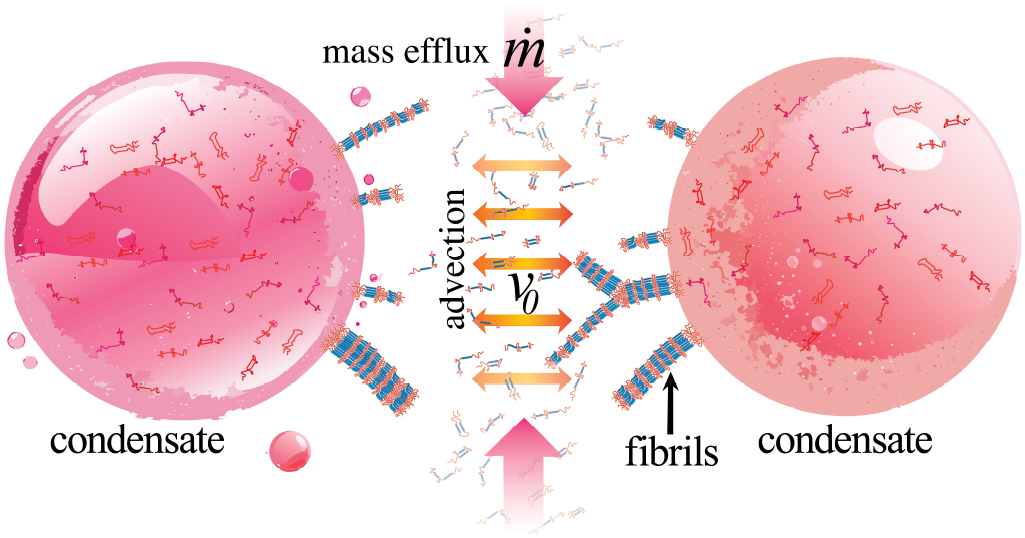
Schematic overview of condensate-to-fibril transition. Rigid, *β*-prone segments (blue) preferentially align and promote fibril formation at the condensate interface, where reduced dimensionality lowers entropic barriers to ordered assembly. Sustained non-equilibrium molecular influx 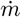 and initial advection velocity *v*_0_ from the dilute phase supply material for fibril elongation and contribute to the transition from a liquid-like to a fibril-dominated, solid-like state.

## II. RESULTS

We model protein chains as coarse-grained polymers containing rigid, *β*-prone segments connected by flexible linkers, driven toward condensate interfaces under controlled molecular influx (see Methods for full details).

### A. Fibrillar nucleation and growth at condensate surfaces

FD-MD simulations show that condensate interfaces favor fibrillization when polymers contain localized rigid, *β*-prone segments with attractive interactions (Fig. 2A), whereas fully flexible chains or chains with randomly distributed rigid segments do not produce fibrillar structures under the same conditions (Fig. 2B,C). In these driven simulations, interfacial growth is sustained by continuous recruitment of chains from the dilute phase: polymers supplied at a rate 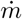 accumulate at the interface and are incorporated into surface-localized assemblies (Fig. 1). At early times, we observe the formation of small interfacial clusters enriched in aligned rigid segments (Fig. 2A). At the interface, the entropic cost of alignment is reduced relative to the bulk, and favorable rigid–rigid attractions stabilize these nascent ordered patches. These clusters serve as protofilament nuclei that template further incorporation of incoming chains. As the simulation proceeds, protofilaments thicken and elongate into surface-anchored fibrillar protrusions that remain anchored to the condensate surface and extend outward into the dilute phase (Fig. 2A).

**FIG. 2.**
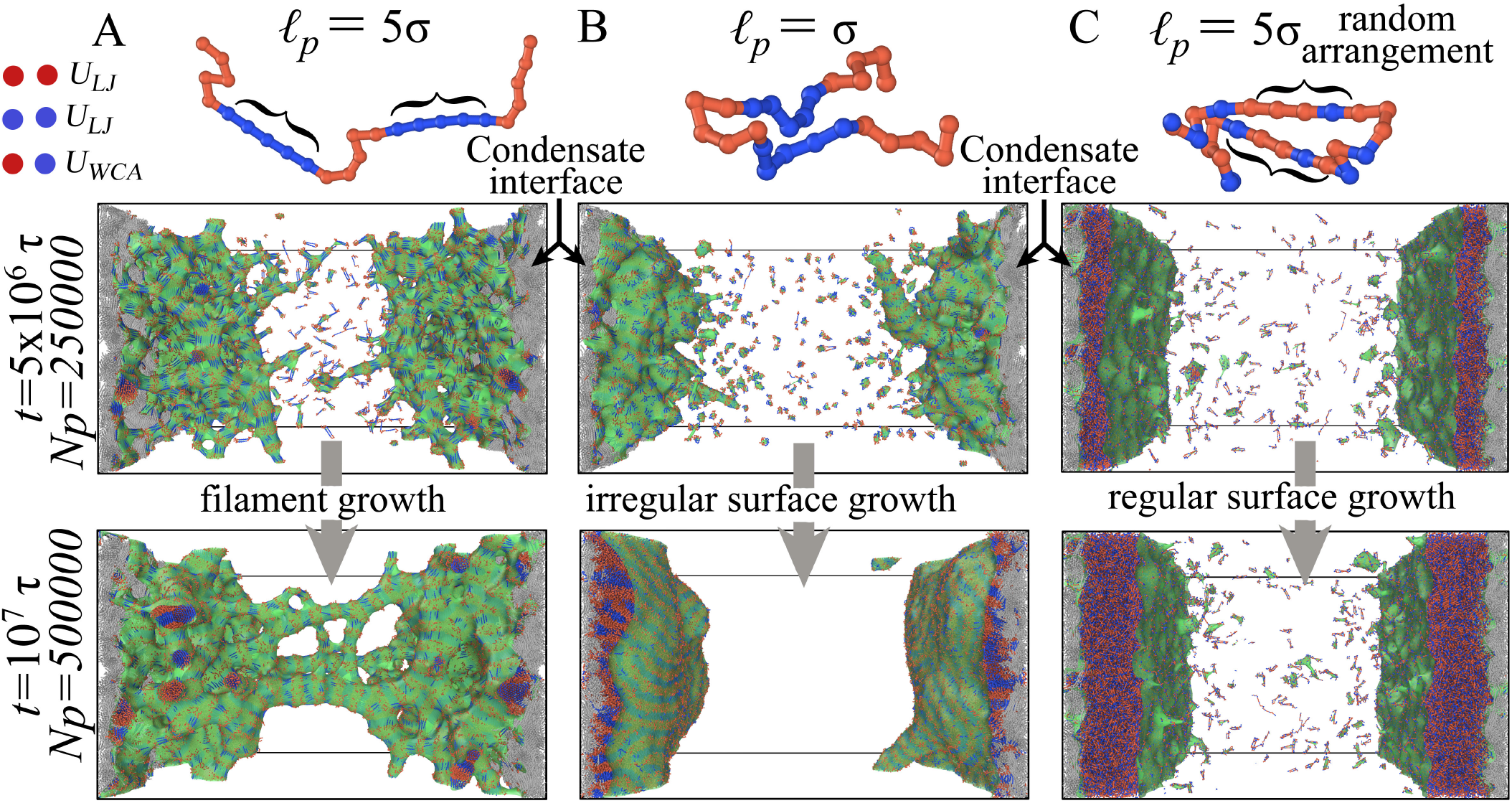
Simulation snapshots of filament growth at biomolecular condensate interfaces. CGMD simulations illustrating the role of sequence-encoded rigidity in fibril formation. (A) Polymers containing two rigid, *β*-prone segments (blue; persistence length *𝓁*_*p*_ = 5*σ*) connected by flexible segments (red) nucleate at the condensate interface and progressively assemble into surface-anchored fibrillar protrusions. With time, these filaments elongate, bundle, and form mesh-like structures that can bridge opposing interfaces. (B) When chains are fully flexible (no rigid *β*-prone segments), interfacial aggregation persists, but no fibrillar structures emerge. (C) Polymers with randomly distributed rigid and flexible segments, despite the presence of a rigid *β*-sheet-like interaction, fail to form filaments and instead promote the growth of extended planar interfaces. These results indicate that both the presence and spatial organization of rigid segments are required in this model for fibril nucleation and growth at condensate interfaces.

The preferential alignment of rigid segments at the interface can be understood through a simple entropic argument. In the bulk dilute phase, a rigid segment of n monomers freely explores all orientations over a solid angle of 4π, so the probability of two such segments spontaneously co-aligning is proportional to (4π)^−(*n*−1)^. At a flat condensate surface, however, geometric exclusion by the dense condensate phase restricts the accessible orientational space to approximately a hemisphere (2π), halving the effective orientational volume and biasing segments toward lying parallel to the interface. This restriction lowers the entropic cost of mutual alignment by ΔΔ*S*≈ (*n*−1) *k*_B_ ln 2 per segment pair relative to the bulk. For a *β*-prone segment comprising *n*≈ 5 beads, this yields ΔΔ*S* ≈4 *k*_B_ ln 2≈ 2.8 *k*_B_, a substantial reduction that lowers the free energy barrier to nucleus formation by several thermal energy units. This geometric entropic effect is the coarse-grained analogue of surface-promoted nucleation in classical nucleation theory [29, 41, 42], and it explains why ordered clusters nucleate preferentially at condensate boundaries rather than in the homogeneous bulk. Extending this argument, the ratio of surface to bulk nucleation rates scales as

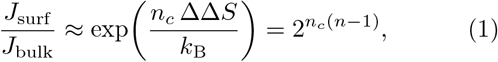

where *n*_*c*_ is the critical nucleus size in units of chains. For *n*_*c*_ = 2 chains of *n* = 5 rigid beads, Eq. (1) gives *J*_surf_ /*J*_bulk_ ≈2^8^ = 256. For the parameter regime explored here, this scaling estimate suggests that orientational entropy alone can enhance interfacial nucleation by roughly two orders of magnitude, providing an order-of-magnitude rationale for why the interface is kinetically favorable as a nucleation site in the model.

We note that pinning of rigid segments at the interface is cooperative: once a segment aligns and makes cohesive contacts with neighbors, each subsequent chain arriving at the interface encounters a partially ordered template that further stabilizes alignment. This cooperative templating is expected to reduce the critical nucleus size at the interface relative to the bulk, providing a molecular basis for the lower nucleation barrier beyond the orientational entropy argument alone. As shown below (Sec. II D), rigid segments that are templated by the interface exhibit strongly suppressed lateral mobility, in contrast to flexible chains which remain diffusively mobile on the surface.

These observations support an interface-promoted mechanism for fibril growth in which (i) interfacial alignment stabilizes critical nuclei and (ii) sustained non-equilibrium recruitment supplies material for directional elongation. To map the conditions under which these morphologies emerge, we constructed a non-equilibrium phase diagram as a function of polymer stiffness (quantified by persistence length *𝓁*_*p*_) and supply rate 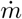 (Fig. 3). Four regimes of surface and fibril growth are observed. For highly flexible chains (*𝓁*_*p*_ ∼ 1), deposition produces isotropic interfacial thickening without directional fibril formation (Fig. 2B), reminiscent of isotropic surface growth (see blue region in Fig. 3). At low but finite stiffness, disordered surface deposits accumulate without long-range order. Above a stiffness threshold, surface-localized amyloid-like fibrils nucleate and elongate robustly. At sufficiently high supply rate and high stiffness, fibrils further bundle and form bridges between opposing interfaces, yielding networked morphologies. In this networked regime, we observe a qualitative subdivision between more fluid-like networks at lower stiffness [43] and mechanically stiff, load-bearing networks at higher stiffness.

**FIG. 3.**
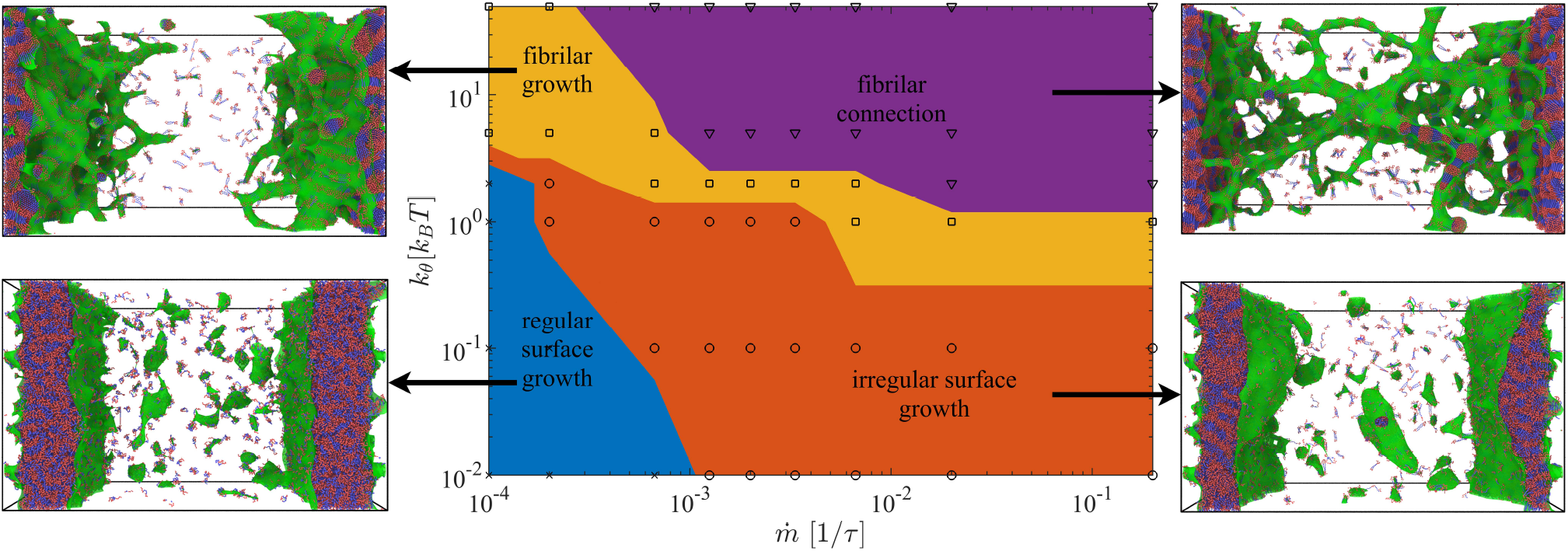
Non-equilibrium phase diagram of interfacial growth morphologies. Phase diagram obtained from FD-MD simulations under sustained mass flux toward a condensate interface, shown as a function of supply rate 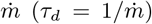 and bending stiffness (or persistence length) of the *β*-prone segment. Four distinct growth regimes are observed: (I) planar surface growth or complete wetting of the condensate surface (fully flexible chains) characterized by smooth interfacial thickening; (II) irregular surface deposition at low but finite stiffness, where disordered aggregates accumulate without long-range order; (III) fibrillar interfacial growth at intermediate-to-high stiffness (partially wetted surface), marked by nucleation and elongation of surface-anchored fibrils; and (IV) bridging and network formation at high stiffness and high supply rate, where fibrils bundle and connect opposing interfaces. Arrows indicate representative snapshots from simulations for each regime.

Critically, varying sequence patterning strongly suppresses fibrillization under the conditions examined, even in the presence of rigid segments (see Fig. 2 C). When *β*-prone motifs are no longer localized along the chain, molecules accumulate broadly at the interface but fail to nucleate fibril formation, producing heterogeneous, craze-like surface morphologies, i.e., disordered, fibril-free deposits with spatially heterogeneous density reminiscent of craze structures observed in polymer fracture mechanics [44]. This finding is consistent with recent experimental evidence that the spatial organization of *β*-prone motifs, not merely their presence, governs the propensity for condensate-to-fibril conversion [45, 46]. The central finding of this section is therefore twofold: localized *β*-prone rigidity functions as a strong determinant of whether condensate interfaces nucleate ordered fibrils or accumulate disordered deposits, and within the present model, varying segment rigidity and molecular supply rate is sufficient to recover four distinct aging morphologies.

### B. Surface Area Growth During Condensate Aging

To characterize non-equilibrium surface growth during protein deposition, we analyzed the morphologies that emerge when aggregates form at condensate interfaces under a sustained molecular flux. Surface growth processes are known to exhibit qualitatively distinct behaviors depending on microscopic growth rules and transport mechanisms [47]. In the present system, we identify three recurring surface growth scenarios: (i) random deposition onto two parallel condensate interfaces, resulting in uniform planar thickening; (ii) localized bundle formation driven by rigid *β*-sheet–forming segments, leading to heterogeneous increases in surface thickness; and (iii) anisotropic fibrillar growth, in which bundles elongate radially from the interface while planar thickening proceeds concurrently.

We find that surface area growth is dominated by the third scenario when rigid segments are present, reflecting the combined effects of intrinsic chain stiffness and reduced post-deposition mobility associated with *β*-sheet content [47]. In contrast, when rigid segments or sequence patterning are absent, growth proceeds exclusively via isotropic planar thickening (scenario i). We therefore analyze surface growth kinetics to distinguish planar deposition from fibril-mediated anisotropic extension.

Figure 4 shows the total surface-associated area as a function of time. At early times, rapid accumulation at the planar interfaces produces a transient regime dominated by surface deposition. The duration of this regime depends on transport conditions: for large drift velocities |*v*_0_|, rapid planar coverage suppresses early fibril formation, whereas smaller drift velocities promote simultaneous fibril nucleation and surface thickening, delaying homogeneous coverage. Cluster volume measurements (Fig. 4 B) corroborate this interpretation. The inset density *ρ*_*n*_(t) further reveals that smaller drift velocities lead to prolonged localization near the interfaces, consistent with enhanced redistribution into fibrillar structures. At long times, once surface coverage is established, a linear growth regime emerges in which surface area increases approximately linearly with time. In this regime, deposited molecules contribute both to continued planar thickening and to radial and longitudinal fibrillar extension. The onset of this linear regime occurs earlier for larger drift velocities, reflecting deposition-dominated dynamics.

**FIG. 4.**
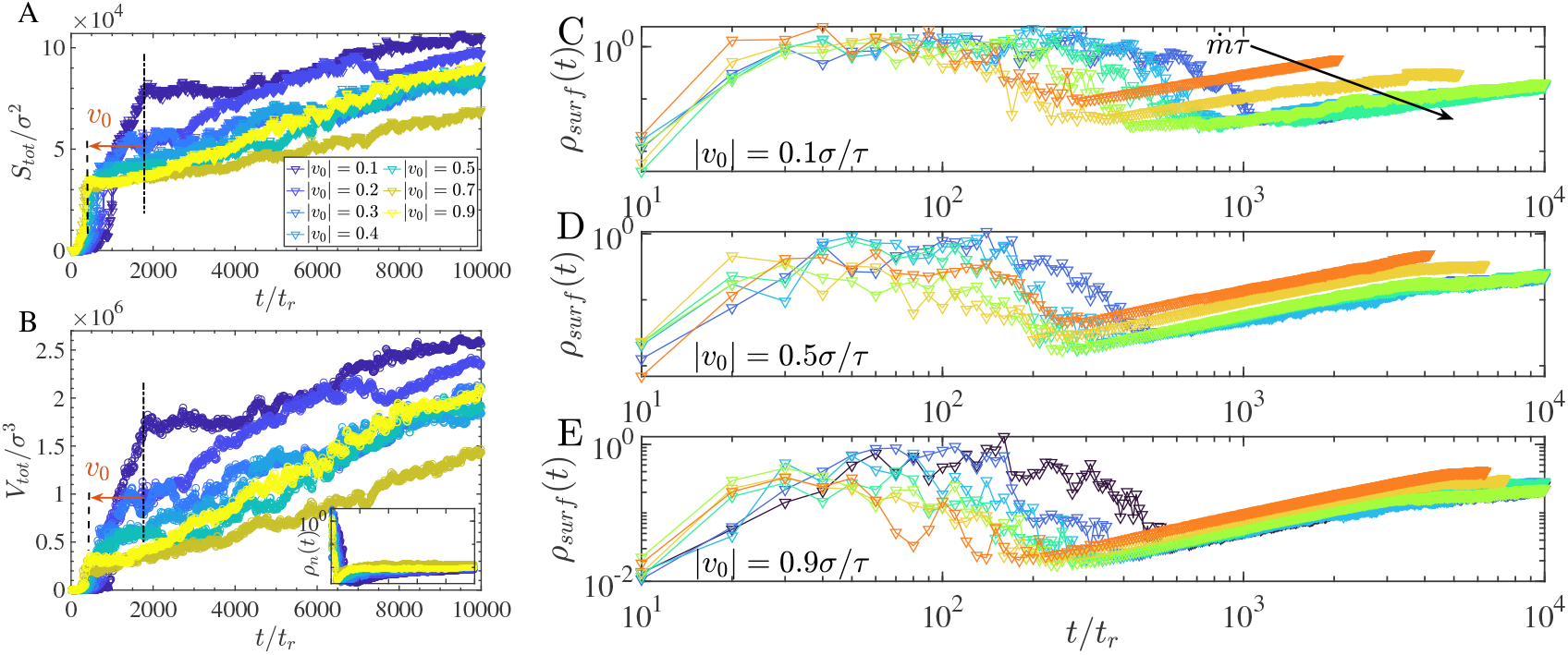
A. Total surface area of molecules deposited onto the condensate interfaces. At early times, exponential growth arises from rapid molecular deposition onto the two parallel surfaces (indicated by vertical lines). For larger drift velocities |*v*_0_|, planar surface formation dominates. For smaller drift velocities, planar deposition and fibril formation occur simultaneously, leading to delayed surface homogenization. Once a continuous surface layer forms, surface area increases approximately linearly due to fibrillar growth. B. The corresponding cluster volume exhibits similar trends. The inset shows the number density *ρ*_*n*_(*t*), indicating that early growth is localized near the condensate surfaces, while later redistribution reflects fibril formation. C-E. Surface density *ρ*_surf_ as a function of time for supply rates 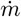 at drift velocities |*v*_0_| = 0.1, 0.5, 0.9 *σ/τ*. At low |*v*_0_| (synthesis-limited regime, C), increasing 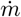 accelerates fibril nucleation onset, advancing the *ρ*_surf_ ∼ *t* scaling regime. At high |*v*_0_| (transport-limited regime, D-E), curves for different 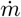 converge while preserving their ordering, indicating saturation of the fibril incorporation rate.

To further distinguish isotropic surface deposition from directional fibrillar growth, we quantified the evolution of the surface density *ρ*_surf_ (*t*) for three drift velocities |*v*_0_| = 0.1, 0.5, and 0.9 *σ/τ* across varying supply rates 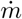 (Fig. 4 C-E). At early times, all curves display a non-monotonic growth–decline–growth trajectory: ρ_surf_ rises sharply, dips by one to two orders of magnitude as the nucleation burst drives a rapid expansion of S(t), then recovers as fibrillar elongation resumes linear surface growth. An analytical explanation of this three-phase trajectory, based on the saturating growth kinetics developed below, is given in Supporting Information (Sec. S1.5 and Fig. S1). At longer times, all cases converge to a scaling regime characterized by *ρ*_surf_ ∼*t*.

At low |*v*_0_| (Fig. 4 C), increasing 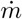 strongly accelerates fibril nucleation onset: higher molecular supply initiates fibrillar growth before the planar wetting layer is fully established, advancing the onset of the *ρ*_surf_ ∼t scaling regime and increasing the overall rate of fibrillar mass accumulation at the interface. This reflects a synthesis-limited regime in which the supply rate 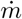 is the dominant kinetic control over fibrillar aging.

At high |*v*_0_| (Fig. 4 D-E), curves for different 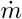 converge while preserving their ordering, indicating a crossover to a transport-limited regime in which chains arrive at the interface faster than the fibril incorporation machinery can process them. In this regime, increasing 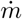 further has a diminishing effect on fibril growth rate; the system is transport-saturated. Together, these two regimes identify a crossover transport rate 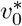 that separates supply-driven from transport-driven fibril growth. To rationalize the two limiting behaviors within a compact expression, we write a hyperbolic saturating function for the effective fibril growth rate,

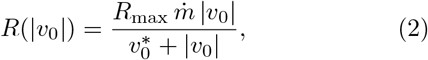

where *R*_max_ is the maximum growth rate at transport saturation, 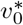 is the half-saturation drift velocity at which the growth rate reaches half its maximum value, and 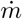 enters as a prefactor encoding molecular supply. In the synthesis-limited limit 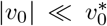, Eq. (2) gives 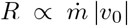, consistent with the strong 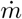 -dependence observed at low drift velocity (Fig. 4C). For 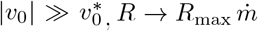, recovering the transport-saturated regime where curves for different 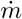 converge (Fig. 4D-E). This hyperbolic form is the simplest rational function consistent with linear growth at small |*v*_0_| and saturation at large |*v*_0_| ; it arises generically whenever a rate-limited incorporation step competes with a flux-limited supply step (see Supporting Information, Sec. S1.2 for a derivation from steady-state mass balance). We note that 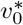 is not a single universal constant but shifts with 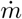 : at low molecular supply, transport saturates fibril incorporation even at modest drift velocities, whereas at high supply, transport remains rate-limiting across the full range of |*v*_0_| explored. Full derivations and the connection to the three-phase *ρ*_surf_ (t) kinetics are provided in Supporting Information (Secs. S1.2–S1.5).

The spatial redistribution underlying these growth modes is further quantified by the inter-interface density profile *ρ*_*N*_ (*x*): at low drift velocity and high supply rate, density progressively migrates from the interfaces toward the central region, confirming fibrillar bridge formation, whereas at high drift velocity it remains interface-localized, consistent with planar thickening (see Supporting Information).

These results reveal that condensate aging is not a single pathway but a family of trajectories selected by molecular supply. At low flux, condensates age slowly through fibril-dominated growth; at high flux, rapid planar saturation suppresses fibrillization [32, 40]. The crossover between these synthesis-limited and transport-limited regimes is characterized by a half-saturation velocity 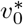 that, while not universal (it shifts with 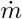), provides a compact organizing parameter for the observed kinetic regimes.

### C. Filament Growth Dynamics

While randomized sequences produce isotropic surface deposition with quasi-steady total area *S*(*t*), sequences containing rigid segments exhibit pronounced anisotropic growth. Surface deposition initiates symmetry breaking at localized nucleation sites, from which filamentous protrusions elongate into the dilute phase. Continued molecular recruitment drives both longitudinal extension and radial thickening of these filaments. The emergence of such high-curvature structures leads to a monotonic increase in the total surface area *S*(*t*) with time, in contrast to the quasi-steady behavior observed for randomized sequences. As detailed below and in Fig. 5, the rate of longitudinal elongation depends inversely on drift velocity, reflecting a competition between directed tip recruitment and bulk surface saturation.

**FIG. 5.**
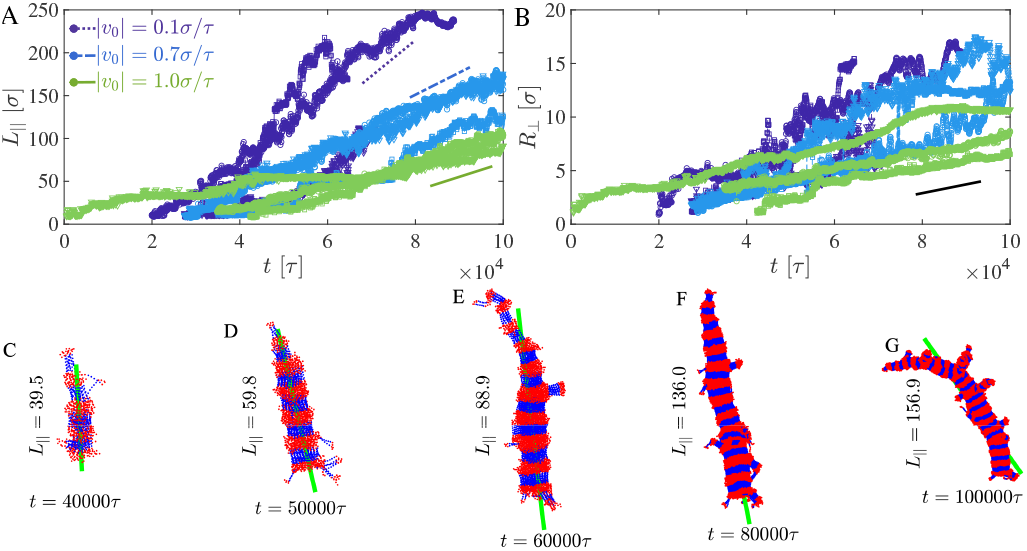
Filament growth kinetics. (A) Longitudinal filament length *L*_‖_ versus time for different drift velocities |*v*_0_|, showing linear elongation with slopes that decrease with increasing |*v*_0_|. (B) Radial extent *R*_⊥_ versus time, showing weak dependence on |*v*_0_|. (C-G) Representative filament configurations at increasing times; green line indicates the principal filament axis.

To quantitatively characterize filament morphology, we computed the gyration tensor of each identified filament cluster,

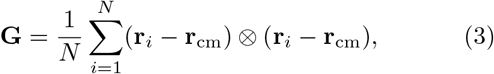

where *N* is the number of beads in the filament, **r**_*i*_ are bead positions, and **r**_cm_ is the filament center of mass. Diagonalization of **G** yields eigenvalues λ_1_ ≥λ_2_≥ λ_3_ and eigenvectors **e**_1_, **e**_2_, **e**_3_, where **e**_1_ defines the fila-ment axis. A characteristic longitudinal length may be estimated as 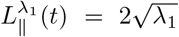 in practice, we define the longitudinal filament length as the end-to-end distance *L*_∥_(*t*). The transverse extent is quantified as 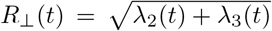, separating axial growth from lateral thickening. The degree of shape anisotropy is further quantified using

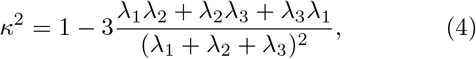

where *κ*^2^→ 1 corresponds to highly elongated structures. We observe *κ*^2^(*t*) ≈ 1 coincident with the onset of filament growth, indicating persistent symmetry breaking during elongation (Fig. 5C–G).

Figure 5A shows that the longitudinal filament length *L*_∥_(t) increases approximately linearly with time,

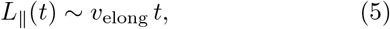

where *v*_elong_ is the effective elongation velocity.

Notably, faster drift velocity slows fibril elongation. For a drift velocity |*v*_0_| = 0.1, we measure *v*_elong_ ≈ 6.0×10^−3^, but as the drift velocity increases to |*v*_0_| = 0.7 and 1.0, the elongation rate decreases to 2.5 ×10^−3^ and 1.5 × 10^−3^, respectively, a four-fold reduction over the full range. This inverse dependence is described by a tip-saturation model,

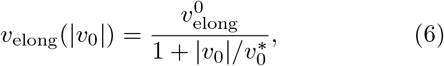

where 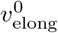 is the elongation rate at vanishing drift and 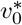 is the crossover velocity at which the tip-docking step becomes rate-limiting. Fitting Eq. (6) to the three mea-sured rates yields 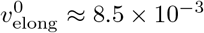 and 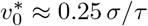,indicating that tip-docking becomes the bottleneck at modest drift velocities. We note that fitting two parameters to three measured rates provides consistency with, but not a stringent test of, the proposed functional form. Mechanistically, this inverse dependence reflects a competition between planar surface coverage and directed tip recruitment: at high drift velocities, incoming chains rapidly saturate the flat interface before anisotropic symmetry breaking can be established, diverting material away from filament tips and slowing elongation. At lower drift velocities, the interface is not prematurely saturated and a greater fraction of arriving chains are captured at the growing filament tips, sustaining faster directional elongation. The derivation of Eq. (6) and its role in the late-time recovery of *ρ*_surf_ are detailed in Supporting Information (Secs. S1.3 and S1.5).

In contrast, the radial growth *R*_⊥_(t) (Fig. 5B) exhibits weak sensitivity to |*v*_0_|, with a characteristic slope of approximately 7.5 ×10^−5^. This suggests that lateral accretion onto filament surfaces is governed primarily by local surface attachment kinetics rather than by global influx dynamics.

The separation between longitudinal and radial growth modes is consistent with two distinct kinetic processes: (i) tip-mediated elongation, controlled by directional molecular recruitment and sensitive to drift velocity, and (ii) surface accretion, driven by local adsorption and largely independent of global transport rates. The linear scaling of *L*_∥_(*t*) is a hallmark of reaction-limited growth, whereas the weak dependence of *R*_⊥_(*t*) on |*v*_0_| is consistent with diffusion-limited lateral attachment. We present this reaction-limited/diffusion-limited classification as the minimal interpretation consistent with the observed scaling, while noting that more detailed mechanistic tests would be needed to rule out alternative explanations.

These two regimes can be distinguished by a simple rate argument. For tip-mediated elongation, an incoming chain must diffuse to the filament tip and adopt a geometrically compatible orientation before it can be incorporated. When this orientational activation step is slower than diffusive arrival, the growth rate is governed by an Arrhenius-like attachment rate,

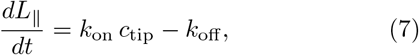

where *c*_tip_ is the local chain concentration at the tip and *k*_on_ encodes the Boltzmann factor for successful orientational docking. At steady state under constant influx, *c*_tip_ is approximately constant, giving *L*_∥_ ∼ *t*, the hallmark of interface-limited (reaction-limited) growth familiar from crystal elongation kinetics [15, 41, 48]. This picture also explains the inverse dependence of *v*_elong_ on |*v*_0_| : at high drift velocity, chains arrive at the tip faster than they can be correctly oriented and incorporated, creating a kinetic bottleneck at the attachment site and reducing the net elongation rate.

In contrast, lateral accretion requires no specific chain orientation: any chain diffusing to the filament surface can adsorb. The rate would then be limited by the flux of chains reaching the cylindrical surface rather than by any activation barrier, consistent with diffusion-limited radial growth. Because local surface diffusion and adsorption, not the global drift velocity |*v*_0_|, control this process, *R*_⊥_ is insensitive to drift velocity, consistent with what we observe. The anisotropic eigenvalue spectrum (λ_1_≫λ_2_≈ λ_3_) confirms persistent filament symmetry breaking throughout growth.

The key observation from this analysis is that fibril elongation and lateral thickening respond oppositely to molecular transport conditions, consistent with distinct rate-limiting steps for the two growth modes. If this interpretation holds, it implies that the aspect ratio of fibrillar products is not fixed by sequence alone but is tunable through transport: slower delivery would favor longer, thinner fibrils, while faster delivery would yield shorter, thicker deposits [25, 49].

### D. Dynamic Arrest During Surface-Mediated Fibril Growth

The preceding sections establish that rigid *β*-prone segments drive anisotropic fibrillar growth at condensate interfaces, with surface area expansion and filament elongation characterized by distinct kinetic regimes. Recent observations show that crowding-induced dynamical heterogeneity in IDP assemblies leads to anomalous, non-Fickian diffusion [50]. A complementary molecular-level signature of this structural transition is the progressive loss of individual chain mobility as molecules are incorporated into ordered assemblies. To quantify this transition, we computed the mean-squared displacement (MSD) of individual polymer chains, defined as ⟨Δ*r*^2^(*t*)⟩ ∼ *t*^*α*^, where the exponent *α* characterizes the underlying transport mechanism. MSDs were calculated for the center-of-mass trajectories of individual chains from the time of their introduction into the simulation box, and ensemble averages were taken over all deposited molecules.

At early times, newly introduced chains exhibit superdiffusive, ballistic motion due to the imposed drift toward the condensate interfaces, characterized by an effective exponent *α* ≈ 2 (Fig. 6A). As deposition proceeds and interfacial aggregates begin to form, molecular motion slows markedly. In systems that undergo fibrillar growth, the MSD exhibits a pronounced crossover to sub-diffusive dynamics, with an effective exponent *α* ≈ 0.1, eventually approaching a plateau indicative of dynamic arrest. We note that MSD measurements in this non-equilibrium setting involve averaging over molecules introduced at different times and deposited at varying interfacial heights. Consequently, ensemble averaging may yield nontrivial effective exponents. Nevertheless, a key observation is that once chains are incorporated into surface-bound aggregates or fibrillar structures, their mobility becomes strongly constrained, leading to subdiffusive behavior. The observed exponent *α* ≈ 0.1 is consistent with diffusion in highly constrained, viscoelastic, or caged environments, where motion is restricted by neighboring aligned segments within growing fibrils. This behavior is analogous to cage dynamics observed in glass-forming systems and dense polymer networks [51], and is consistent with the viscoelastic aging and dynamical slowdown reported in protein condensates [12, 52]. The crossover from ballistic to subdiffusive motion thus reflects the progressive immobilization of chains as they become incorporated into protofilaments and mature fibrillar assemblies.

**FIG. 6.**
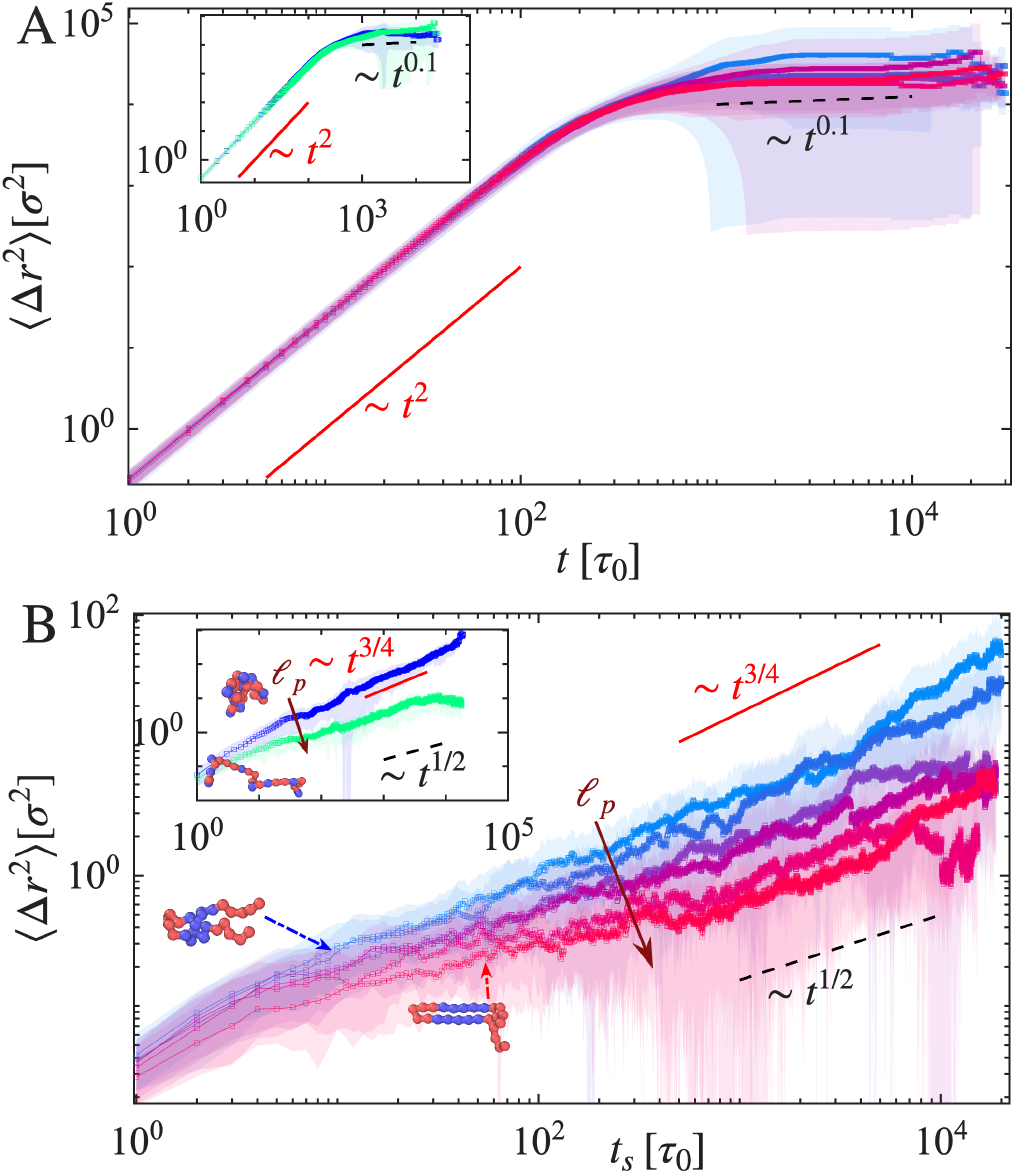
Mean-squared displacement (MSD) of polymer chains. (A) MSD from time of introduction into the system, showing ballistic transport at early times followed by a plateau indicative of dynamical arrest. Color gradient (blue to red) represents increasing *𝓁*_*p*_ at fixed drift velocity and mass influx. Inset: randomized sequence controls. (B) MSD after surface adsorption, showing subdiffusive scaling that strengthens with increasing *𝓁*_*p*_. Inset: randomized sequence controls exhibit similar trends.

To disentangle early-time transport from interfacial dynamics, we introduce a shifted time origin *t*_*s*_, defined after molecules are fully incorporated into the surface. MSDs computed relative to *t*_*s*_ (Fig. 6B) reveal clear distinctions between systems with different chain rigidities. Chains with persistence length 𝓁_*p*_ = 5*σ* exhibit stronger subdiffusive behavior, scaling approximately as ⟨Δ*r*^2^(*t*)⟩ ∼ *t*^1*/*2^, whereas flexible chains display weaker subdiffusion with *α* ≈ 3/4. These trends are consistent with randomized sequence controls (Fig. 6B, inset), which exhibit similar scaling behavior. Importantly, systems that do not undergo fibrillization, such as those with randomized sequence architectures lacking localized rigid *β*-prone motifs, retain diffusive or weakly subdiffusive dynamics throughout deposition, with *α* ≈ 3/4. In these cases, surface-adsorbed chains maintain significant lateral mobility, indicating the absence of structural constraints necessary to induce dynamic arrest.

The transition in MSD scaling thus provides a molecular-level counterpart to the macroscopic growth regimes identified above: the reaction-limited tip elongation and diffusion-limited lateral accretion described in the preceding section are consistent with progressive kinetic trapping of individual chains within the growing fibrillar scaffold. Although a fraction of molecules remains diffusive in the bulk at intermediate times, continued non-equilibrium recruitment ultimately leads to their incorporation and arrest at the interface during fibril elongation. Within our simulations, the onset of subdiffusive dynamics thus serves as a useful simulation-level marker of fibrillar maturation, distinguishing systems that undergo structural ordering from those that remain liquid-like. Rigid *β*-prone segments not only template ordered structures but progressively trap molecules within them, suggesting that MSD scaling may provide an accessible readout of the extent of fibrillar arrest at condensate interfaces.

## III. DISCUSSION

Within our coarse-grained model, two minimal ingredients, sequence-encoded *β*-prone rigidity and nonequilibrium molecular influx, are sufficient to produce condensate aging pathways ranging from liquid-like thickening to persistent fibrillar arrest, as observed in recent experiments [22, 33, 35, 37, 45]. By varying only segment stiffness and molecular supply rate in FD-MD simulations, we mapped a non-equilibrium phase diagram containing four qualitatively distinct interfacial morphologies within this framework: uniform wetting, disordered deposition, surface-anchored fibrillar growth, and inter-condensate bridging networks. Within the present model, this morphological richness emerges primarily from two control parameters, suggesting that the diversity of aging phenotypes observed across different protein systems may in part reflect variations in a lowdimensional parameter space rather than fundamentally different mechanisms.

At the mesoscale, distinct aggregation regimes give rise to characteristic statistical signatures. Flexible sequences lacking interacting *β*-sheet–forming segments produce smooth, uniformly thickened interfaces (uniform wetting), whereas rigid, *β*-prone segments induce symmetry breaking and the formation of anisotropic, columnar fibrillar protrusions. These structural transitions are captured by anisotropic surface growth statistics, including surface density scaling (*ρ*_surf_ ∼ *t*), cluster shape anisotropy (*κ*^2^→ 1), and the onset of dynamic arrest in the MSD, providing experimentally accessible markers of condensate maturation. Three minimal analytical models connect the simulation observables to broader theory: a surface-enhanced nucleation rate (Eq. 1) that estimates the geometric entropic advantage of the interface over the bulk; a hyperbolic saturating growth law (Eq. 2) that unifies the synthesis-limited and transport-limited regimes under a single expression; and a tip-saturation model for elongation velocity (Eq. 6) that rationalizes the inverse dependence of *v*_elong_ on drift velocity. Together, these expressions reduce the multidimensional FD-MD phase space to a compact set of physically interpretable parameters. By comparing driven and undriven MD conditions, we find that fibrillization can arise from collective aggregation in dilute regions even in the absence of an imposed drift. However, the formation of a continuous bridge between dilute-phase fibrils and the condensate interface requires a sustained influx of molecules near the interface. The key advantage of FD-MD is not only its ability to capture interface-driven fibrillization, but also to provide quantitative control over flux parameters, enabling systematic mapping of rate-dependent morphological regimes. In particular, it reveals the kinetic competition between planar surface saturation and directed fibril elongation, phenomena that remain inaccessible to equilibrium simulations. We note that a complementary nonequilibrium flux-driven approach has recently been applied to model directed molecular transport through condensates [40], underscoring the broader utility of driven simulation frameworks for condensate biology.

Several limitations of the present framework should be noted. First, the planar slab geometry used here provides computational tractability and well-defined interfacial normals, but it eliminates curvature-dependent effects present at real droplet surfaces, including Laplace pressure gradients and curvature-driven molecular sorting, that may modulate nucleation barriers and fibril growth directionality at spherical condensate interfaces. Second, the isotropic Lennard-Jones attraction combined with angular stiffness is a defensible coarse-grained proxy for *β*-sheet assembly, but it does not capture the directionality of backbone hydrogen bonding, strand registry, or the cooperative nature of *β*-sheet elongation. These features could influence the detailed morphology and kinetics of fibril growth beyond what the present model resolves. Third, the analytical scaling estimates (Eqs. 1, 2, 6) are minimal models intended to rationalize limiting regimes rather than to provide globally quantitative fits. Future work incorporating curved interfaces, explicit directional interactions [53, 54], and direct nucleation rate measurements will be needed to assess the quantitative transferability of these findings to specific protein systems.

In sum, our results suggest that condensate aging can be decomposed into two independently controllable axes: sequence-encoded rigidity governs the structural competence for fibril formation, while molecular flux selects which morphological pathway is taken. If the separation between these two axes holds in vivo, it raises a notable possibility: strategies that modulate molecular supply (e.g., protein expression or intracellular transport rates) could in principle redirect aging trajectories without altering the protein sequence itself [49, 55]. More broadly, FD-MD provides a general framework for connecting molecular sequence features to driven mesoscale organization, applicable beyond amyloid systems to any condensate whose aging is shaped by the interplay of sequence-encoded structure and non-equilibrium mass transport.

## IV. METHODS

### A. Molecular model of condensate and fibril-forming chains

Protein chains were modeled as coarse-grained polymers composed of alternating rigid, *β*-prone segments and flexible linker regions. This architecture captures key features of proteins such as hnRNPA1 and Tau, which contain localized fibril-prone motifs embedded within intrinsically disordered regions. Rigid segments encode a propensity for ordered *β*-sheet-like alignment, while flexible linkers represent disordered spacers that regulate conformational entropy, interfacial alignment, and recruitment dynamics.

Fibril-forming biomolecular chains are represented as beads connected by finitely extensible nonlinear elastic (FENE) bonds with potential

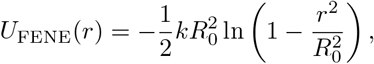

where *r* is the distance between adjacent beads. We used a spring constant *k* = 30 *ε/σ*^2^ and a maximum extensibility *R*_0_ = 1.5*σ*, with bead diameter *σ* defining the unit of length.

Backbone rigidity within *β*-prone segments is imposed via a harmonic angular potential,

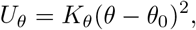

where *θ* is the angle between consecutive bonds, *θ*_0_ = 180^°^, and *K*_*θ*_ = 50 ε/*k*_B_*T*. To probe the role of chain stiffness, *K*_*θ*_ was systematically varied across simulations.

Nonbonded interactions are modeled using a Lennard-Jones (LJ) potential,

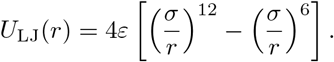

Excluded-volume interactions were enforced by truncating the potential at *r*_*c*_ = 2^1*/*6^*σ*. Attractive interactions between rigid segments were captured by extending the cutoff to *r*_*c*_ = 4*σ*, allowing for cohesive aggregation while using isotropic pairwise interactions.

The use of an isotropic potential to model *β*-sheet-like inter-segment attraction warrants justification. Real *β*-sheet interactions are inherently directional, arising from backbone hydrogen bonds spaced ≈ 4.7 Å along the fibril axis and ≈ 10 Å between strands. However, in our coarse-grained representation each bead encompasses multiple residues, so the directional character of individual hydrogen bonds is averaged over the relevant length scales. The effective segment–segment interaction therefore acquires an approximately isotropic character whose range and depth encode the collective attraction of multiple backbone contacts, consistent with established coarse-grained protein and amyloid models [42, 48, 56]. Critically, the angular stiffness potential *U*_*θ*_ independently enforces the extended, rod-like geometry that is a prerequisite for *β*-strand compatibility: only segments that are simultaneously rigid and in close lateral proximity interact attractively through the extended LJ cutoff. The two terms thus cooperate to reproduce the essential anisotropy of *β*-sheet assembly, geometric compatibility encoded in *U*_*θ*_ and cohesive inter-strand attraction encoded in *U*_LJ_, without requiring explicit directional hydrogen-bond potentials.

Individual chains consist of 15–55 beads and contain one to five rigid *β*-prone segments embedded within flexible linkers. To isolate the role of sequence organization, control simulations were performed in which rigid segments were either removed or randomly distributed along the chain. These controls enabled direct assessment of the necessity of spatially organized rigidity for interfacial nucleation and fibrillar growth.

### B. Simulation protocol

Molecular dynamics simulations were performed using the LAMMPS package [57]. Equations of motion were integrated using a velocity-Verlet scheme with a timestep of 0.005 *τ* in reduced Lennard–Jones units, where *ε* = 1, *σ* = 1, and particle mass m = 1. These units define the characteristic time 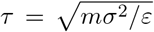, temperature *T* ^*^ = *ε*/*k*_B_, and density *ρ*^*^ = *σ*^−3^. To model fibril growth between extended condensate interfaces, two planar condensate slabs (acting as part of two condensate surfaces) were positioned at opposite boundaries along the x-axis (non-periodic) of a simulation box with dimensions *L*_*x*_ = 200*σ*, with periodic dimensions *L*_*y*_ = *L*_*z*_ = 120*σ*. The condensate slab consisted of approximately 2 × 10^5^ beads, forming stable, viscoelastic interfaces. Between the slabs, a total of *N* = 20,000 polymer chains were introduced at random positions at time intervals *τ*_*d*_ to represent dilute-phase molecules.

### C. Flux Implementation

To model sustained molecular recruitment to condensate interfaces, we implement a thermostatted driven-insertion protocol in which polymer chains are continuously introduced into the simulation box at a controlled rate. Because chains are added over time, the total particle number *N* increases throughout the simulation; the system is therefore an open, non-equilibrium setup with constant volume and temperature maintained by a Nosé– Hoover thermostat. In standard equilibrium molecular dynamics, transport toward the interface is solely diffusion-driven, leading to slow, spatially heterogeneous aggregation in the dilute phase. In particular, polymers may nucleate and form branched aggregates before reaching the condensate surface, preventing controlled surface-mediated growth.

To mimic the experimental situation in *in vitro* systems [21, 22, 32], where a chemical potential gradient drives molecular transport toward a condensate acting as a sink, we introduce a directed influx of polymers. This is achieved by inserting chains at the center of the simulation box with an initial velocity bias toward the condensate interfaces. Specifically, newly introduced polymers are assigned velocities 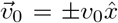, such that half of the chains move toward the +*x* interface and the other half toward the −*x* interface, inserted into the system at rate 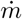 (chains per unit time).

FD-MD thus introduces two independently controllable non-equilibrium parameters with distinct physical meanings. The drift velocity |*v*_*0*_| governs how rapidly individual chains traverse the dilute phase toward the interface, analogous to molecular mobility or cytoplasmic flow rate. The supply rate 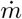 (chains per unit time) sets the total rate of molecular delivery, the quantity most directly analogous to protein synthesis or expression rate as controlled by experimentalists. The deposition timescale 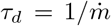 is used interchangeably with 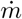 inwhat follows. These two parameters have separable effects on interfacial growth morphology: |*v*_0_| controls how chains reach the interface, while 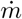 controls how many chains are available to do so (see SI Fig. 2 & Fig. 3). The subsequent dynamics are evolved under the thermostat at constant temperature, ensuring that thermal fluctuations are maintained while preserving the net directional bias at early times. Importantly, the magnitude of *v*_0_ is chosen to be sufficiently small that polymer conformations and interfacial structure remain indistinguishable from those obtained under purely diffusive conditions, while significantly accelerating the rate of interface encounter. We note that under purely diffusive, thermostatted conditions (no imposed drift, *v*_0_ = 0), polymers may nucleate branched aggregates in the bulk dilute phase before reaching the condensate surface, preventing controlled surface-mediated growth. The directed insertion protocol suppresses this competing pathway by biasing chains toward the interface before bulk aggregation can occur.

### D. Surface growth analysis

To quantify non-equilibrium surface growth in FD-MD simulations, we identified surface-associated molecules based on their proximity to the condensate interfaces. A polymer chain was classified as surface-associated if at least one bead was located within a cutoff distance *r*_*c*_ = 2*σ* from either planar condensate surface. Variations of *r*_*c*_ within the range 1.5*σ*–3*σ* did not qualitatively affect the results.

The total surface-associated area was computed by identifying connected clusters of surface-associated beads and estimating their exposed interfacial area, capturing contributions from both planar surface coverage and protruding fibrillar structures. In parallel, the total cluster volume was computed as the sum of bead volumes within the largest surface-associated cluster, allowing discrimination between lateral spreading and volumetric growth. Surface coverage kinetics were quantified using the surface density *ρ*_surf_ (*t*) = *N* _surf_ (*t*)/*A* _int_, where *N*_surf_ (*t*) is the number of surface-associated beads and *A*_int_ is the nom-inal area of a single planar condensate interface.

To characterize spatial redistribution and interfacial bridging, we computed the number density profile of the largest surface-associated cluster along the interface separation axis, 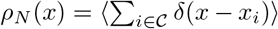, where 𝒞 denotes beads belonging to the largest surface-associated cluster. Accumulation of density in the central region between interfaces was used as an indicator of fibrillar bridge formation.

## Data Availability

All simulation input files, analysis scripts, and data necessary to reproduce the results are available at [repository URL to be added upon acceptance].

## Acknowledgment

This work was supported by funds from the National Institute of General Medical Sciences, with grant no. R35 GM138243 awarded to D.A.P.

