## Supporting Information for "*β*-motifs and molecular flux promote amyloid nucleation at condensate interfaces"

### Supporting Information: $\beta$-motifs and molecular flux promote amyloid nucleation at condensate interfaces

Subhadip Biswas[1*] and Davit A Potoyan[1,2†]
[1]*Department of Chemistry, Iowa State University, Ames, IA 50011, USA and*
[2]*Department of Biochemistry, Biophysics and Molecular Biology, Iowa State University, Ames, IA, 50011, USA*
(Dated: April 9, 2026)

#### S1. ANALYTICAL ESTIMATES FOR INTERFACIAL NUCLEATION AND GROWTH KINETICS

In this section we provide minimal kinetic and statistical-mechanical derivations that rationalize the scaling forms used in the main text. These derivations are not intended to be exact, but rather to capture the dominant physical mechanisms governing interfacial nucleation and flux-driven growth.

##### S1.1. Surface-enhanced nucleation rate

We begin by estimating the ratio of nucleation rates in the bulk and at the condensate interface. In classical nucleation theory, the nucleation rate is written as

$$J \sim J_0 \exp\left(-\frac{\Delta G^\ddagger}{k_\mathrm{B} T}\right), \tag{S1}$$

where $\Delta G^\ddagger$ is the free energy barrier and $J_0$ is a kinetic prefactor. We write the barrier as $\Delta G^\ddagger = \Delta H^\ddagger - T\Delta S^\ddagger$. Assuming that the dominant difference between bulk and surface nucleation arises from the entropic term, we write

$$\Delta G^\ddagger_\mathrm{surf} - \Delta G^\ddagger_\mathrm{bulk} \approx -T\left(\Delta S^\ddagger_\mathrm{surf} - \Delta S^\ddagger_\mathrm{bulk}\right) = -T\Delta\Delta S^\ddagger. \tag{S2}$$

Neglecting differences in prefactors, the ratio of nucleation rates becomes

$$\frac{J_\mathrm{surf}}{J_\mathrm{bulk}} \approx \exp\left(\frac{\Delta\Delta S^\ddagger}{k_\mathrm{B}}\right). \tag{S3}$$

To estimate $\Delta\Delta S^\ddagger$, we consider the orientational entropy of a rigid segment composed of $n$ beads. Formation of a nucleus requires approximate co-alignment of these segments. In the bulk, each orientational degree of freedom explores a solid angle of $4\pi$, so the accessible orientational phase space scales as

$$\Omega_\mathrm{bulk} \sim (4\pi)^{n-1}. \tag{S4}$$

At a flat interface, steric exclusion by the dense phase restricts orientations to approximately a half-space, giving

$$\Omega_\mathrm{surf} \sim (2\pi)^{n-1}. \tag{S5}$$

The reduction in orientational entropy per segment is therefore

$$\Delta\Delta S_\mathrm{seg} = k_\mathrm{B} \ln\frac{\Omega_\mathrm{bulk}}{\Omega_\mathrm{surf}} = (n-1)k_\mathrm{B}\ln 2. \tag{S6}$$

*
†

For a critical nucleus composed of $n_c$ chains, we approximate

$$\Delta\Delta S^{\ddagger} \approx n_c(n-1)k_{\mathrm{B}} \ln 2. \tag{S7}$$

Substituting into the rate expression yields

$$\frac{J_{\mathrm{surf}}}{J_{\mathrm{bulk}}} \approx \exp\left(n_c(n-1)\ln 2\right) = 2^{n_c(n-1)}. \tag{S8}$$

This estimate neglects correlations along the polymer backbone and the finite softness of the interface, and should therefore be interpreted as a scaling-level argument.

##### S1.2. Effective growth rate under flux

We next derive a minimal expression for the effective growth rate of interfacial aggregates under sustained molecular influx. Let $c_I$ denote the surface-associated concentration of chains near the interface. The evolution of $c_I$ is governed by a balance between arrival and incorporation,

$$\frac{dc_I}{dt} = J_{\mathrm{in}} - J_{\mathrm{inc}}. \tag{S9}$$

We model the incoming flux as

$$J_{\mathrm{in}} \sim a\,\dot{m}\,|v_0|, \tag{S10}$$

where $\dot{m}$ is the molecular supply rate, $|v_0|$ is the drift velocity, and $a$ is a geometric factor relating bulk influx to interfacial arrival. Incorporation into ordered structures is assumed to be saturable due to a finite number of effective growth sites. We therefore write

$$J_{\mathrm{inc}} = \frac{k\,c_I}{K + c_I}, \tag{S11}$$

where $k$ is the maximal incorporation rate and $K$ is a characteristic concentration scale at which the incorporation rate reaches half its maximum. At steady state, $J_{\mathrm{in}} = J_{\mathrm{inc}} = R$, where $R$ is the effective growth rate. This saturable form arises generically whenever a finite number of incorporation sites process arriving molecules: at low interfacial concentration the rate is proportional to $c_I$, while at high concentration the sites are fully occupied and the rate plateaus. Rather than solving explicitly for $c_I$, we construct a minimal interpolation consistent with the limiting regimes. For small drift velocity, arrival is rate-limiting and

$$R \propto \dot{m}|v_0|. \tag{S12}$$

For large drift velocity, arrival is no longer limiting and the rate saturates at a value proportional to the supply,

$$R \to R_{\mathrm{max}}\dot{m}. \tag{S13}$$

The simplest rational function consistent with these limits is the hyperbolic saturating form

$$R(|v_0|) = \frac{R_{\mathrm{max}}\,\dot{m}\,|v_0|}{v_0^* + |v_0|}, \tag{S14}$$

where $v_0^*$ is a half-saturation velocity marking the crossover between transport-limited and transport-saturated regimes. This expression is the minimal interpolation between the two limiting kinetic regimes: linear growth at low drift and saturating growth at high drift. It arises from the steady-state balance between a flux-limited supply and a site-limited incorporation step, without requiring any specific enzymatic or catalytic mechanism.

##### S1.3. Tip-saturation model for filament elongation

We now consider the elongation of fibrillar structures at the interface. Let incoming chains arriving at the interface be partitioned between two pathways: incorporation at filament tips and adsorption onto already-covered planar surface regions. Denoting the corresponding effective rates as $k_{\text{tip}}$ and $k_{\text{surf}}$, the fraction of chains contributing to tip growth is

$$f_{\text{tip}} = \frac{k_{\text{tip}}}{k_{\text{tip}} + k_{\text{surf}}}. \tag{S15}$$

We assume that $k_{\text{tip}}$ is approximately constant, while $k_{\text{surf}}$ increases with drift velocity due to more rapid planar surface coverage,

$$k_{\text{surf}} = \alpha |v_0|, \tag{S16}$$

where $\alpha$ is a proportionality constant. Substituting gives

$$f_{\text{tip}} = \frac{k_{\text{tip}}}{k_{\text{tip}} + \alpha |v_0|} = \frac{1}{1 + |v_0|/v_0^*}, \tag{S17}$$

where

$$v_0^* = \frac{k_{\text{tip}}}{\alpha}. \tag{S18}$$

If each successful tip incorporation event contributes a mean length increment and the baseline elongation rate at vanishing drift is $v_{\text{elong}}^0$, the effective elongation velocity is

$$v_{\text{elong}}(|v_0|) = v_{\text{elong}}^0 \, f_{\text{tip}} = \frac{v_{\text{elong}}^0}{1 + |v_0|/v_0^*}. \tag{S19}$$

##### S1.4. Linear growth of filament length

Finally, we justify the observed linear growth of filament length. Let $N_{\text{tip}}(t)$ denote the number of successful incorporation events at filament tips. If each event increases the filament length by a characteristic increment $\delta\ell$, then

$$L_{\parallel}(t) = L_0 + \delta\ell \, N_{\text{tip}}(t). \tag{S20}$$

Under steady influx conditions, the rate of successful incorporation is approximately constant,

$$\frac{dN_{\text{tip}}}{dt} = k_{\text{eff}}. \tag{S21}$$

It follows that

$$\frac{dL_{\parallel}}{dt} = \delta\ell \, k_{\text{eff}} \equiv v_{\text{elong}}, \tag{S22}$$

and therefore

$$L_{\parallel}(t) = L_0 + v_{\text{elong}} t. \tag{S23}$$

This result is characteristic of reaction-limited growth under steady supply conditions.

##### S1.5. Surface density dynamics: three-phase kinetics and non-monotonic growth

The cluster surface area $S(t)$ and the accumulated molecular mass $\dot{m}\,t$ evolve on different timescales during fibril formation. Their ratio defines the surface density

$$\rho_{\mathrm{surf}}(t) = \frac{\dot{m}\,t}{S(t)}, \tag{S24}$$

which measures how tightly mass is packed onto the growing surface. FD-MD simulations show that $\rho_{\mathrm{surf}}(t)$ is non-monotonic: it rises sharply, dips by one to two orders of magnitude, then recovers. This behavior follows directly from the three-phase kinetics captured by Eqs. (S14) and (S19).

*a. Phase 1 — nucleation expansion.* Before a stable fibrillar nucleus forms, the cluster surface grows through the accumulation and lateral ordering of adsorbed chains. The rate of surface expansion is proportional to the existing surface area (autocatalytic) and is driven by the arrival flux governed by Eq. (S14),

$$\frac{dS}{dt} = R_{\mathrm{nucl}}(|v_0|)\,S, \qquad R_{\mathrm{nucl}} = \frac{\alpha_0\,|v_0|}{v_0^* + |v_0|}, \tag{S25}$$

giving exponential surface growth $S(t) = S_0\,e^{R_{\mathrm{nucl}}\,t}$. Substituting into Eq. (S24):

$$\rho_{\mathrm{surf}}(t) = \frac{\dot{m}}{S_0}\,t\,e^{-R_{\mathrm{nucl}}\,t}. \tag{S26}$$

This $t\,e^{-R_{\mathrm{nucl}}\,t}$ form rises, peaks at

$$t_{\mathrm{peak}} = \frac{1}{R_{\mathrm{nucl}}} = \frac{v_0^* + |v_0|}{\alpha_0\,|v_0|}, \tag{S27}$$

then declines exponentially as $S$ grows faster than the linearly accumulating mass. Higher drift velocity increases $R_{\mathrm{nucl}}$ and therefore shifts the peak to earlier times while reducing its amplitude, consistent with the simulation data (Fig. S1a).

*b. Phase 2 — fibrillar elongation.* At $t = t^*$, a nucleation burst drives $S$ to a much larger value $S^* \gg S_0$. After this transition the dominant growth mechanism is fibril tip elongation, which is limited by tip saturation according to Eq. (S19). The surface then grows linearly,

$$S(t) = S^* + v_{\mathrm{elong}}(|v_0|)\,(t - t^*), \qquad t > t^*, \tag{S28}$$

and the surface density recovers,

$$\rho_{\mathrm{surf}}(t) = \frac{\dot{m}\,t}{S^* + v_{\mathrm{elong}}(|v_0|)\,(t - t^*)}. \tag{S29}$$

For $t \gg t^*$ this approaches the asymptote $\dot{m}/v_{\mathrm{elong}}(|v_0|)$, which *increases* with $|v_0|$ because tip saturation (Eq. (S19)) reduces $v_{\mathrm{elong}}$ at higher drift — the opposite trend from Phase 1.

*c. Origin of the deep dip.* The sharp minimum in $\rho_{\mathrm{surf}}$ near $t = t^*$ arises from the large ratio $S^*/S_0 \sim 10^2$: the nucleation burst creates $\sim 100\times$ more surface area almost instantaneously, while the accumulated mass at that moment has grown by only $t^*/t_{\mathrm{peak}} \sim 5\times$. The depth of the dip is therefore set by the geometry of nucleation, not by transport.

*d. Saturating kinetics controls both phases.* Both the peak position (Eq. (S27), through $R_{\mathrm{nucl}}$) and the recovery rate (Eq. (S29), through $v_{\mathrm{elong}}$) are governed by the same half-saturation velocity $v_0^*$. This unification means the entire non-monotonic trajectory of $\rho_{\mathrm{surf}}$ is captured by Eqs. (S14) and (S19) with a single kinetic parameter.

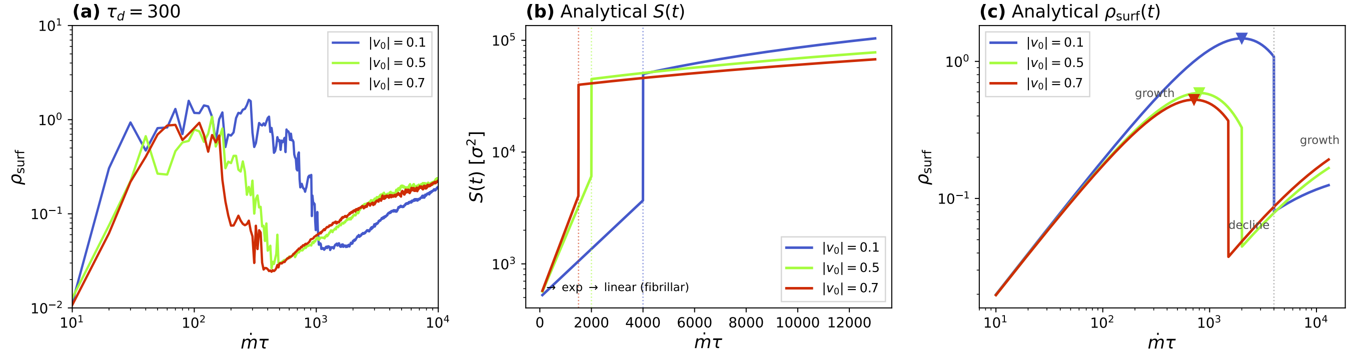


FIG. S1. **Surface density dynamics from FD-MD simulations and the minimal saturating-kinetics analytical model.** (a) $\rho_{\rm surf}(\dot{m}\tau)$ at fixed synthesis rate $\tau_d = 300$ for drift velocities $|v_0| = 0.1$ (blue), 0.5 (green), and $0.7\,\sigma/\tau$ (orange). Higher $|v_0|$ shifts the peak to earlier times and lowers its amplitude, consistent with Eq. (S27). (b) Analytical three-phase model for $S(t)$: exponential expansion during nucleation (Phase 1) transitions to linear fibrillar elongation (Phase 2) at $t^*$ (dotted vertical lines); higher $|v_0|$ accelerates the transition but slows the subsequent elongation via tip saturation (Eq. (S19)). (c) Analytical $\rho_{\rm surf}(t) = \dot{m}\,t/S(t)$ from the three-phase model (Eqs. (S26)–(S29)). The growth–decline–growth shape arises naturally: exponential $S$ growth in Phase 1 suppresses $\rho_{\rm surf}$ after the initial peak (triangles), while linear $S$ growth in Phase 2 allows recovery. Annotations mark the three kinetic regimes for the $|v_0| = 0.1$ trajectory.

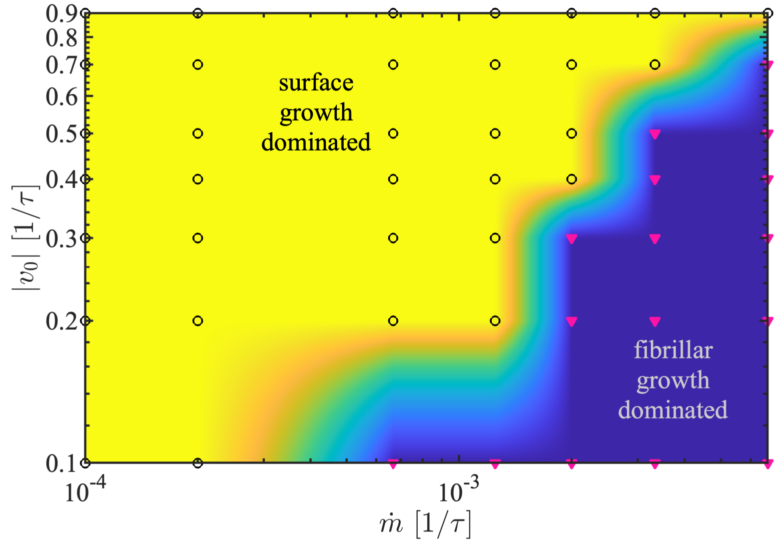


FIG. S2. Phase diagram of whether the system is fibrillar growth dominated or surface growth dominated after depositing 20000 molecules with different initial drift $v_0$ and 0.9, and molecule influx rate $\dot{m}$.

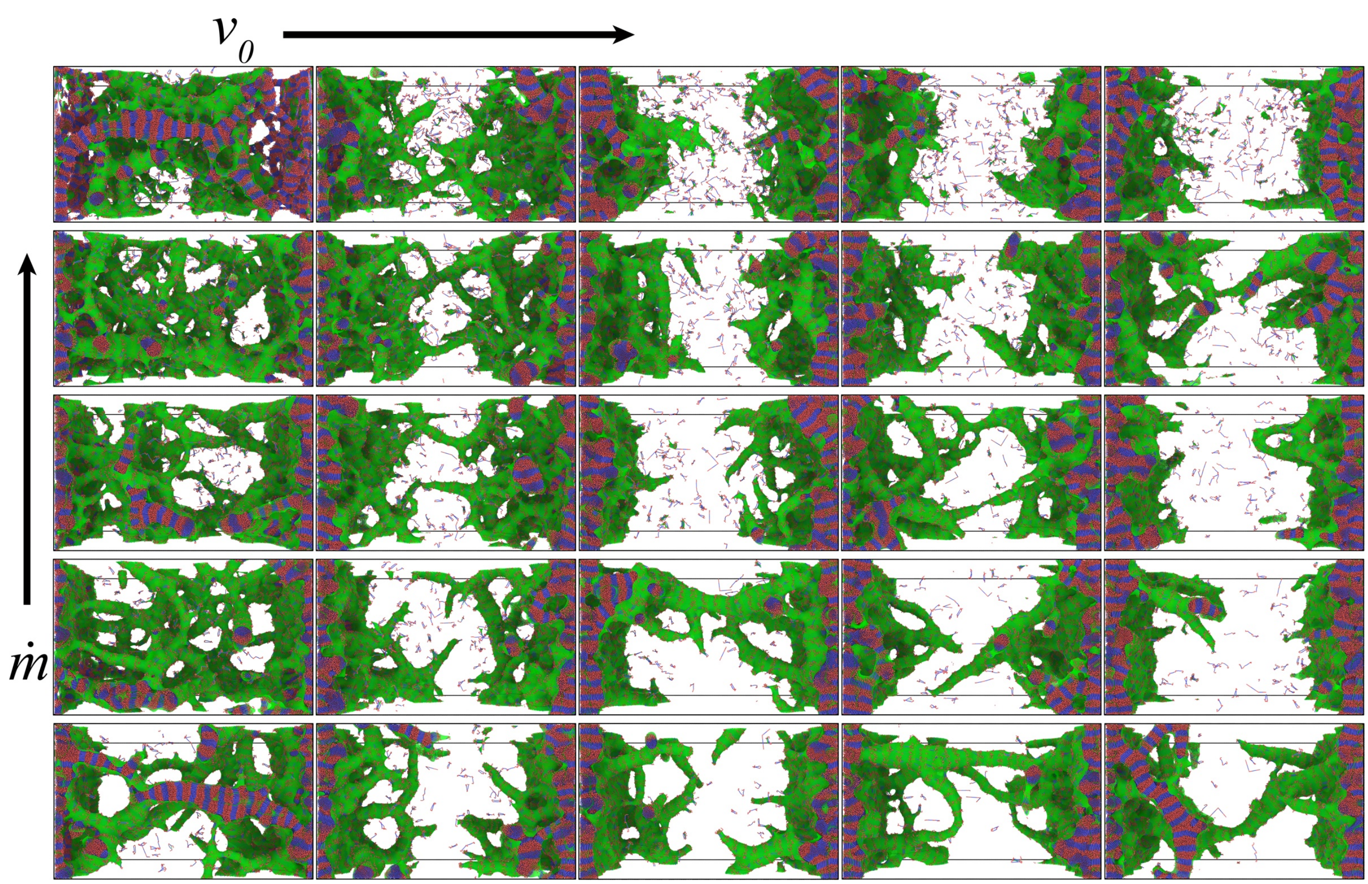


FIG. S3. Phase diagram of simulation snapshots of end frame after depositing 20000 molecules with different initial drift $v_0 = 0.1, 0.2, 0.4, 0.5$ and 0.9, where molecule influx rate $\dot{m}$ goes 1/1.5, 1/3, 1/5, 1/8, 1/15 keeping $\ell_p = 5\sigma$ constant.
